# Mol-AE: Auto-Encoder Based Molecular Representation Learning With 3D Cloze Test Objective

**DOI:** 10.1101/2024.04.13.589331

**Authors:** Junwei Yang, Kangjie Zheng, Siyu Long, Zaiqing Nie, Ming Zhang, Xinyu Dai, Wei-Ying Ma, Hao Zhou

**Affiliations:** School of Computer Science, National Key Laboratory for Multimedia Information Processing, Peking University-Anker Embodied AI Lab, Peking University; School of Artificial Intelligence, National Key Laboratory for Novel Software Technology, Nanjing University; Institute for AI Industry Research (AIR), Tsinghua University; PharMolix Inc.

## Abstract

3D molecular representation learning has gained tremendous interest and achieved promising performance in various downstream tasks. A series of recent approaches follow a prevalent framework: an encoder-only model coupled with a coordinate denoising objective. However, through a series of analytical experiments, we prove that the encoderonly model with coordinate denoising objective exhibits inconsistency between pre-training and downstream objectives, as well as issues with disrupted atomic identifiers. To address these two issues, we propose Mol-AE for molecular representation learning, an auto-encoder model using positional encoding as atomic identifiers. We also propose a new training objective named 3D Cloze Test to make the model learn better atom spatial relationships from real molecular substructures. Empirical results demonstrate that Mol-AE achieves a large margin performance gain compared to the current state-of-the-art 3D molecular modeling approach. The source codes of Mol-AE are publicly available at https://github.com/yjwtheonly/MolAE.

## 1. Introduction

Pre-training based molecular representation learning has shown remarkable performance across various molecular understanding tasks, such as drug discovery (Pinzi & Rastelli, 2019; Adelusi et al., 2022), molecular property prediction (Luo et al., 2022; Liu et al., 2022b; Zhou et al., 2023; Yu et al., 2023; Ju et al., 2023) and reaction prediction (Gastegger et al., 2021; Schwaller et al., 2021). Early approaches tend to model 1D SMILES (Wang et al., 2019; Guo et al., 2021; Honda et al., 2019) or 2D graphs (Li et al., 2021; Lu et al., 2021; Fang et al., 2022b; Xia et al., 2022). More recently, there has been a growing interest in 3D molecular data, with its inclusion of 3D structure information providing more comprehensive information of molecules. Consequently, numerous studies have explored the individual or joint pre-training of 3D modality for better molecular understanding (Liu et al., 2022a; Fang et al., 2022a; Zhou et al., 2023; Luo et al., 2022; Jiao et al., 2023; Yu et al., 2023; Feng et al., 2023)(please refer to Appendix A for detailed related works).

In 3D molecular representation learning, two core techniques are prevalently adopted: (i) **Encoder-only model**. The latent representations output by the encoder are directly used for learning both pre-training tasks and downstream tasks. (ii) **Coordinate denoising objective**. This objective introduces random noise to atom coordinates, and the model is trained to reconstruct the original coordinates. For simplicity, we refer to the framework adopting **en**coder-only model and **c**oordinate **d**enoising objective as *EnCD*. Approaches following *EnCD* framework have demonstrated significant efficacy in various 3D molecular understanding tasks and have achieved state-of-the-art performance in certain benchmarks (Luo et al., 2022; Zhou et al., 2023; Yu et al., 2023). However, there are two inherent problems preventing better performance of *EnCD*:

- The encoder-only model struggles to address the inconsistency between pre-training and downstream objectives. Downstream molecular understanding tasks typically require global information, whereas the pre-training of *EnCD* focuses on local coordinate information (van Tilborg et al., 2022; Zhang et al., 2023). This leads to poor transferability of the features learned by the encoder.
- The coordinate denoising may lead to unstable training and introduce unrealistic noise into the model. In 3D molecular representation learning, coordinates serve two roles simultaneously. One role, analogous to the words in texts, represents the content being reconstructed. The other role, analogous to word positions in texts, should remain stable to let the model know which atom is being reconstructed. The twisted optimization caused by these two roles during coordinate denoising makes model training and convergence challenging. Additionally, as many previous works have pointed out (Wang et al., 2022a; Feng et al., 2023), denoising objective may cause the model to learn unreliable noisy distributions, thereby impacting its performance.

To address these two issues, we introduce a novel approach named Mol-AE (Molecular Auto-Encoder), which incorporates two key designs: (i) To mitigate the inconsistency of the encoder-only model arising from objectives, we propose using an auto-encoder model for pre-training and discard decoder for downstream tasks, since we observe that such inconsistency has a more severe impact on deeper layers (Section 3.1). (ii) To tackle the issues associated with coordinate denoising, we propose a novel objective termed the 3D Cloze Test (Figure 1). Instead of disrupting both roles of the coordinates simultaneously, the objective provides additional positional encoding (PE) during disrupting coordinates to enable the model to discern atom identities, thus achieving stable training. At the same time, the objective uses dropping instead of adding noise for disruption, enabling the model to focus only on remaining noise-free substructures.

**Figure 1.**
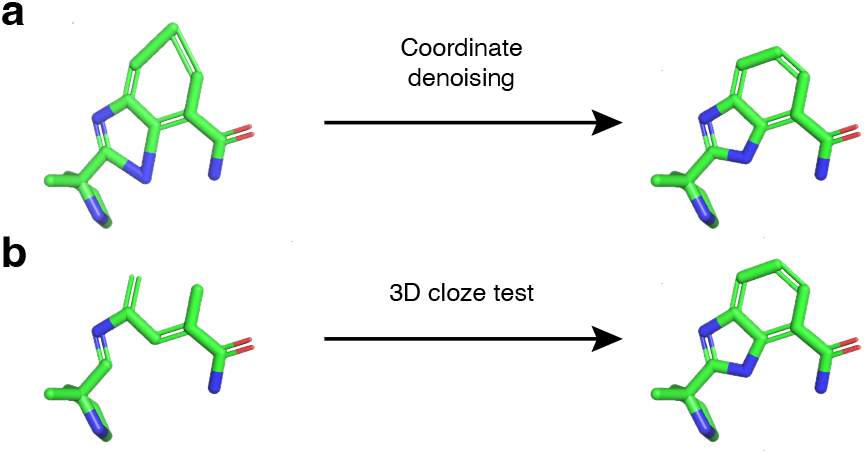
Illustrations of the coordinate denoising (**a**) and 3D cloze test (**b**) objectives.

Extensive experimental results demonstrate that Mol-AE consistently outperforms various molecular representation learning methods across a diverse set of molecular understanding tasks.

Our contributions are summarized as follows:

- We provide an analysis of two inherent problems present in the mainstream frameworks currently used for 3D molecular representation learning and prove the necessity of the auto-encoder architecture and positional encoding in atoms. These problems lack a systematic formulation and analysis in previous works.
- We introduce a straightforward yet powerful model, named Mol-AE, to solve the two problems. Mol-AE employs an auto-encoder architecture as the backbone model and leverages the novel 3D Cloze Test objective.
- Extensive experimental results demonstrate that Mol-AE achieves state-of-the-art performance on a standard molecular benchmark, including various molecular classification and molecular regression tasks.

## 2. Preliminaries

In this section, we will clarify the problem formulation of 3D molecular representation learning, and introduce the most widely-used coordinate denoising method.

### 2.1 Problem Formulation

A 3D molecule ℳ can be seen as a set of *n* atoms, i.e., 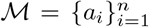 Each atom *a*_*i*_ further consists of its type ***t***_*i*_∈ ℕ and coordinate ***c***_*i*_ ∈ ℝ^3^. We denote the type and coordinates of all the atoms in as **T**∈ ℕ^*n*×1^ and **C**∈ ℝ^*n*×3^, respectively.

Briefly, the goal of 3D molecular representation learning is to train a parameterized encoder *q*_*ϕ*_ to map a molecule to an informative latent representation **Z**∈ ℝ^*n*×*d*^ for downstream tasks, where *d* is the dimension of the latent space. Formally, the objective of 3D molecular representation learning can be expressed as a Kullback–Leibler divergence (KL) term:

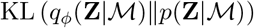

where *p*(**Z** | ℳ) is the real optimal distribution, which characterizes the latent space we desire. However, as *p*(**Z** |ℳ) is usually unknown, we introduce a parameterized decoder *p*_*θ*_ and use the following formula ^1^ as the actual objective:

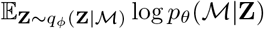

This objective is a classical auto-encoder objective.

### 2.2 Coordinate Denoising

In 3D molecular representation learning, the goal of coordinate denoising is to learn the structural knowledge of 3D molecules while avoiding falling into trivial solutions. Specifically, as the dimension of the latent space is significantly larger than that of the coordinate space, i.e., 3 ≪ *d*, the classical auto-encoder objective may lead to simple identity mappings for both *q*_*ϕ*_ and *p*_*θ*_. In practice, a denoising variant of the objective is used to avoid such trivial solutions.

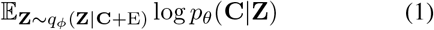

where E∈ ℝ^*n*×3^ is the noise acting on coordinates. Notably, as only partial coordinates are noised, some rows of E contain random noises, while the remaining rows consist of zero vectors. Here, since denoising is only applied to coordinates, we replace ℳ in the auto-encoder objective with the corresponding **C** without loss of generality. Currently, a prevalent choice is using a Transformer encoder as *q*_*ϕ*_ and an SE(3) equivariant head containing simple Multi-Layer Perceptron (MLP) as *p*_*θ*_ (Luo et al., 2022; Zhou et al., 2023; Yu et al., 2023; Jiao et al., 2023).

Two points are worth noting here. First, many related works employ pair-wise distance reconstruction as the training objective, but since this is equivalent to reconstructing SE(3)-invariant coordinates (Satorras et al., 2021), we then exclusively focus on coordinate reconstruction. Second, for models employing an extremely simple decoder, although their mathematical objective aligns with that of an autoencoder, we do not conventionally categorize them as adopting an auto-encoder structure. Since the reconstruction loss directly impacts the latent representation in these cases, we still name these models as encoder-only methods, e.g., BERT (Devlin et al., 2018) and Uni-Mol (Zhou et al., 2023).

## 3. Analysis of *EnCD* Framework

In this section, we will discuss two inherent problems faced by the framework *EnCD* (**En**coder-only model with **C**oordinate **D**enoising objective), which is the current best practice in 3D molecular representation learning and has been utilized in a series of previous works (Luo et al., 2022; Zhou et al., 2023; Yu et al., 2023).

Specifically, (i) encoder-only models cannot handle the inconsistency between pre-training and downstream objectives and (ii) the twisted denoising objective leads to unstable training. Both of them make *EnCD* cannot fully exploit the potential of 3D molecule pre-training.

Here, we employ Uni-Mol (Zhou et al., 2023) and Transformer-M (Luo et al., 2022), two representative *EnCD* methods, along with four widely used molecular property prediction datasets, i.e., BACE, BBBP, HIV, and MUV, to empirically verify these two problems. We will report the results of Uni-Mol here and present the results of Transformer-M in Appendix C, as the conclusions drawn from both methods are consistent. For more dataset details, please refer to Appendix D.

### 3.1. Inconsistencies between Objectives

We will demonstrate the inconsistency between the pretraining objective and the downstream objective in encoder-only molecular representation learning models. Specifically, this inconsistency becomes apparent when the model fails to improve its performance on downstream tasks as its capabilities increase during pre-training.

Following Tenney et al. (2019), we design a layer-wise probing experiment to verify this inconsistency. We first fix the Uni-Mol model and then extract the features from the *L*-th layer, where *L* ∈ {1, *…*, 15 }. Subsequently, we feed the features to an extra prediction head and fine-tune this head with pre-training and downstream tasks, respectively. For pre-training tasks, we use the SE(3)-equivariant heads (Zhou et al., 2023) with the same configuration as the extra prediction heads. For downstream tasks, the extra prediction heads are two-layer MLPs.

As shown in Figure 2a, when using features from the deeper Uni-Mol layer, we can reconstruct 3D molecules, i.e., pretraining task, more accurately. However, as observed from Figure 2b, the performance of most downstream tasks does not improve consistently. In general, the features from the deepest layer do not achieve the best performance and intermediate layer representations often perform better in most cases. Combining the results, we can conclude that **features from deeper layers of encoder-only models exhibit greater performance in pre-training tasks, but this capability does not consistently translate into downstream tasks**.

**Figure 2.**
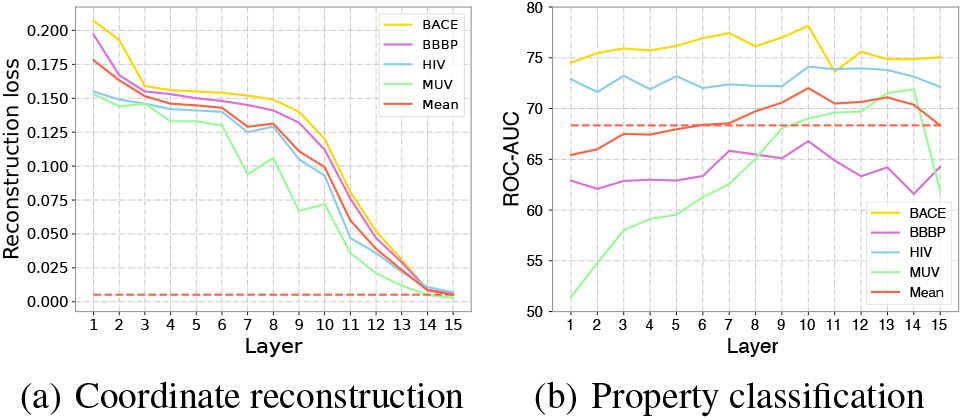
In vanilla Uni-Mol, there are inconsistencies in the modeling capabilities for pre-training and downstream tasks.

Similar phenomena have also been discovered in encoderonly pre-training models for natural language, e.g., BERT and RoBERTa (Devlin et al., 2018; Ethayarajh, 2019; Cai et al., 2020). For example, Tenney et al. (2019) demonstrate that different types of linguistic information are hierarchically represented from shallower to deeper layers of BERT. Both experimental results and related literature motivate us to avoid using a simple encoder-only architecture for 3D molecular representation learning.

### 3.2 Twisted Optimization of *Content* and *Identifier* in Coordinate Denoising

In this section, we will demonstrate that due to the coupling of *Content* and *Identifier*, the coordinate denoising objective for 3D molecular representation learning is actually twisted.

Specifically, there are two types of information, i.e., *Content* and *Identifier*, that play crucial roles in reconstruction-based objectives, e.g., Masked Language Modeling (MLM) for language representation learning and Coordinate Denoising (CD) for 3D molecular representation learning. Both of them are used as the inputs of the objectives, each serving distinct purposes:

- *Content*: For reconstruction-based objectives, *Content* refers to the information that needs to be reconstructed. It should be corrupted in some form so that the model can reconstruct it in order to learn meaningful knowledge.
- *Identifier*: For reconstruction-based objectives, *Identifier* is the anchor for the model to be aware of which part of *Content* needs to be reconstructed. It should be stable and remain uncorrupted.

Here, we use the widely adopted MLM as an example to further elucidate the roles of these two types of information. Specifically, MLM randomly masks partial tokens within a sentence and requires models to reconstruct the masked tokens. Positional encoding is applied to masked tokens as well, helping models accurately reconstruct these tokens by outlining their relationships with unmasked tokens. For MLM objective, the masked tokens are the disrupted *Content* and the positional encoding assigned to masked tokens serves as the *Identifier*. By utilizing these two types of information, MLM efficiently learns linguistic knowledge from natural language, becoming the standard objective for many pre-trained language models (Devlin et al., 2018; Liu et al., 2019b).

On the contrary, due to the coupling between *Content* and *Identifier*, there exists an inherent conflict for the CD objective. Specifically, when applying the CD objective, the coordinates of partial atoms, serving as the *Content*, need to be disrupted by adding random noise. However, these coordinates also serve as the *Identifier* to describe the relationships between the disrupted atoms and the remaining atoms. As previously discussed, they should not be disrupted. The multiple roles of atom coordinates lead to inherent conflicts within the CD, making it a twisted training objective.

We design experiments to further analyze the twisted optimization issue during the coordinate denoising training process. We augment the original Uni-Mol model by adding positional encoding (PE) to each atom to provide stable *Identifier*. Here, the positions of different atoms are determined by the atom order in the SMILES^2^, and this approach is referred to as Uni-Mol-PE. As shown in Figure 3, compared to the original Uni-Mol, the Uni-Mol-PE exhibits lower reconstruction errors, smaller loss fluctuations, and better convergence during training. This indicates that **introducing stable *Identifier* can indeed help the model distinguish between different atoms** to reconstruct the structural information better. Additionally, we can observe that as the disruption intensity increased, Uni-Mol becomes increasingly challenging to train stably. However, with the addition of PE, even in cases of significant disruption (intensity = 5^°^*A*), the model is able to converge stably. But, when evaluating on downstream tasks, we do not observe consistent performance improvement with this straightforward method (Appendix B). We analyse this phenomenon and therefore propose our 3D Cloze Test objective in Section 4.2, which not only promotes pre-training stability but also consistently improves performance on downstream tasks.

**Figure 3.**
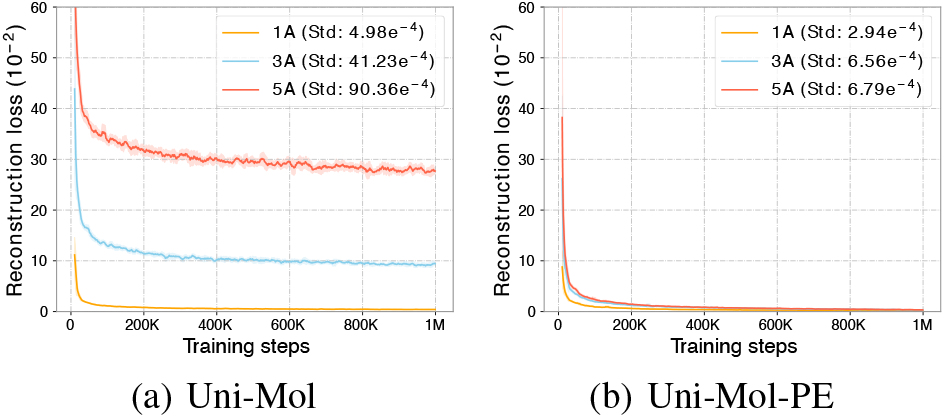
Under different disruption intensities, the impact of introducing PE as an additional Identifier to vanilla Uni-Mol

## 4. 3D Molecular Representation Learning With Mol-AE

In this section, we will detail our 3D molecular representation learning model Mol-AE. As shown in Figure 4, a 3D molecule contains two types of information: the 3D structure and the atom type. Since atom type modeling is a well-defined problem and can be easily achieved by atomic MLM objective (Xia et al., 2022; Wang et al., 2019), thus, we mainly focus on how to model 3D structure and efficiently address the aforehead mentioned problems in *EnCD*.

**Figure 4.**
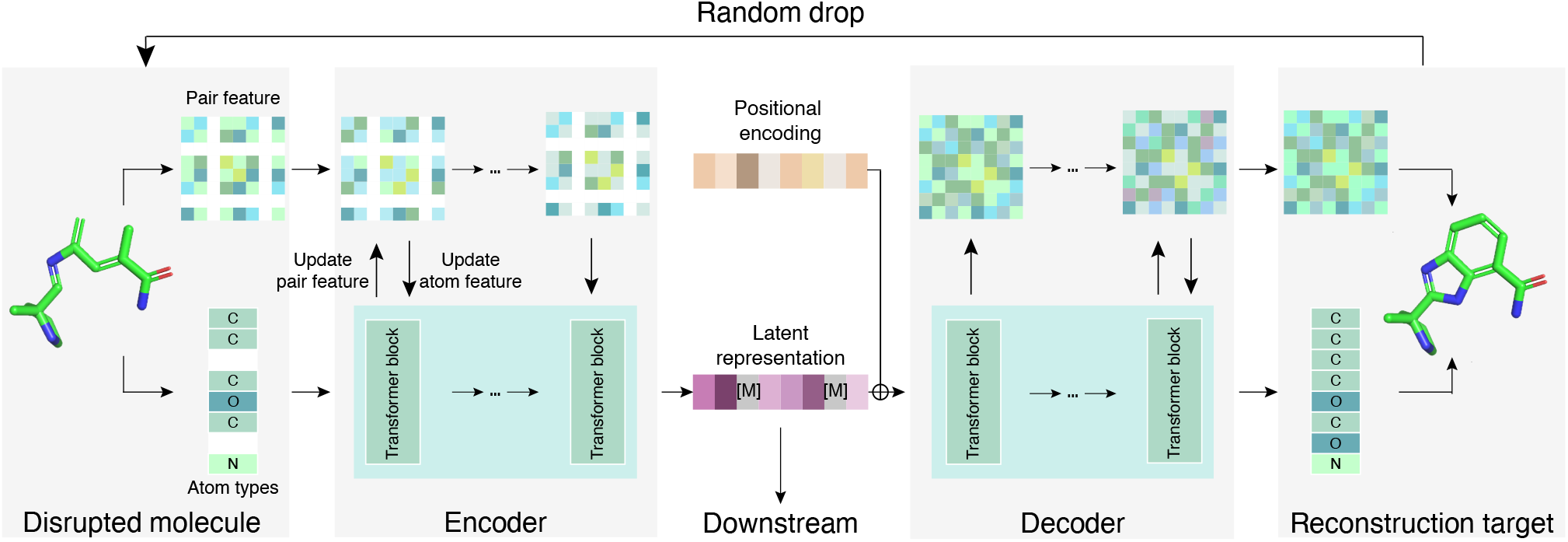
The overall framework of Mol-AE.

In particular, Mol-AE encompasses two fundamental designs: the utilization of an auto-encoder model structure and the incorporation of the 3D Cloze Test objective for model optimization. We will elaborate on our 3D-aware auto-encoder model in Section 4.1, introduce the design of our 3D Cloze Test objective in Section 4.2, and ultimately outline the pre-training and fine-tuning process for Mol-AE in Section 4.3.

### 4.1 3D Information Awared Auto-Encoder

*EnCD* approaches have typically employed a straightforward SE(3) head to instantiate decoder *p*_*θ*_. However, as illustrated in Section 3.1, an excessively simple decoder can lead to a substantial influence of the pre-training objective on the latent representation, thereby impacting its transferability to downstream tasks. Therefore, in Mol-AE, both the *p*_*θ*_ and *q*_*θ*_ utilize the Transformer architecture (Vaswani et al., 2017) as the backbone. This choice is motivated by the Transformer has demonstrated great efficacy in capturing 3D information, as highlighted in recent studies (Luo et al., 2022; Yu et al., 2023; Zhou et al., 2023).

#### Transformer Block

Transformer comprises a series of Transformer blocks, each block contains a multi-head selfattention layer and a feed-forward layer. Denote 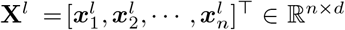 as the input for the *l*-th block, each 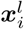 is the *l*-th representation of *a*_*i*_. For any input **X**^*l*^, the **X**^*l*+1^ is generated as follows:

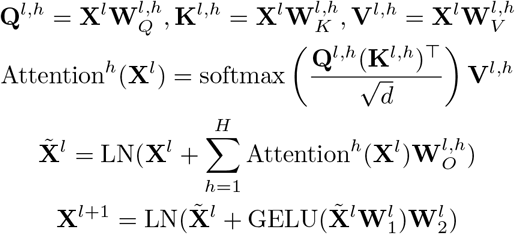

where 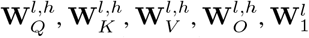 and 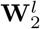 are learnable parameters, *H* is the number of attention heads, and LN is layer normalization (Ba et al., 2016) operation.

#### 3D Aware Pair-Wise Feature

The vanilla Transformer cannot model 3D information, since the coordinates are continuous and invariant under global rotation and translation. Thus, we follow existing approaches (Luo et al., 2022) to encode the Euclidean distance between atom pairs as additional pair-wise features. To be specific, we use the Gaussian Basis Kernel function (Scholkopf et al., 1997) to map the distance between *a*_*i*_ and *a*_*j*_, to a pair feature 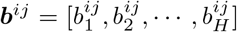 . We denote the pair feature of the *l*-th Transformer layer as ***b***^*ij*,*l*^, and set ***b***^*ij*,0^ = ***b***^*ij*^. To capture better pair-wise relationship, we update ***b***^*ij*,*l*+1^ along with the forward process of Transformer:

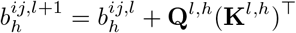

And then we incorporate the edge feature into the vanilla Transformer attention function to enable 3D modeling:

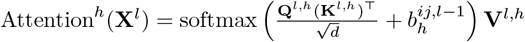

#### Encoder and Decoder in Mol-AE

The encoder *q*_*ϕ*_ comprise *L*^enc^ layers of Transformer blocks. After processing through this encoder, the 3D coordinates information **C** is effectively encoded into 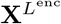, thus we directly set the latent representation 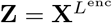. Our decoder *p*_*θ*_ consists of *L*^dec^ layers of Transformer blocks. Please refer to Section 6.3 for empirical study and further discussion about decoder depth. And since 3D structure has already been encoded into **Z**, we initialize the input pair features of the decoder to be all zeros.

### 4.2. Objective of 3D Cloze Test

With the auto-encoder model structure, 3D Cloze Test objective can be formalized as follows:

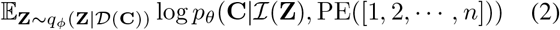

For each 3d coordinate matrix **C**, we first drop a random portion of atoms and corresponding coordinates in drop module 𝒟. And then encode remaining substructures with the transformer-based encoder *q*_*ϕ*_. Because we drop some atoms in the encoder input, we use an insertion operation ℐ to insert the embedding of `[MASK]` token as the representation of dropped atoms into the encoder output **Z**. Then, we add PE on the expanded latent representation ℐ(**Z**) and use decoder *p*_*θ*_ to map them back to 3D coordinates and calculate the reconstruction loss. Compared to the coordinate denoising objective (Equation 1), the 3D Cloze Test objective introduces two key operations: adding PE to the decoder and dropping atoms.

#### Adding PE to the Decoder

As demonstrated in Section 3.2, we successfully addressed the twisted optimization and promoted stable optimization by introducing additional PE to the encoder input as an *Identifier*. However, PE serves not only as the *Identifier* but also introduces order information from SMILES to the model, which could potentially introduce harmful bias. Therefore, mitigating this potential bias when introducing the *Identifier* is crucial. In the autoencoder framework, we notice that the encoder and decoder have distinct roles: the encoder learns a representation of the molecule, while the decoder reconstructs the correct molecular structure. Therefore, a simple method to achieve the aforementioned goal is to add PE only to the decoder, meaning we only add PE to the decoder input ℐ(**Z**). This ensures that the model can better distinguish between different atoms, reconstruct molecular structural information more effectively, and guarantee that the positional information bias does not affect the high-quality molecular representations learned by the encoder.

#### Dropping Atoms

As described in many previous works (Wang et al., 2022a; Feng et al., 2023), denoising objective may cause the model to learn unreliable noisy distributions, thus we disrupt the coordinates by dropping a portion of atoms. Specifically, for the input coordinates **C** ∈ ℝ^*n×*3^, we first randomly remove *k* rows, resulting in 𝒟 (**C**) ∈ ℝ^(*n*−*k*)×3^. Subsequently, we utilize an encoder *q*_*ϕ*_ to obtain its latent representation **Z** ∈ ℝ^(*n*−*k*)× *d*^.

### 4.3 Pre-training and Fine-tuning

#### Pre-training

Since the 3D coordinates are invariant under global rotation and translation, we apply a SE(3)-equivariant head(Zhou et al., 2023) to the output representation of the decoder to calculate the final coordinate reconstruction loss. Previous studies have indicated that, although reconstructing SE(3)-equivariant coordinates and reconstructing pairwise distances are theoretically equivalent, using both in experiments can achieve better results. Therefore, we instruct Mol-AE to simultaneously reconstruct coordinates and pairwise distances to effectively model 3D structure. Additionally, since the atom type can significantly influence molecular properties, we randomly mask some atom types and use a classification head to let Mol-AE predict the ground truth atom types. Following Uni-Mol (Zhou et al., 2023), we introduce a special `[CLS]` atom to represent the entire molecule, with the coordinates of `[CLS]` is the center of all atoms.

#### Fine-tuning

We ignore the decoder and only utilize the encoder for downstream molecular property prediction tasks. During fine-tuning, we refrain from dropping any atoms. We directly add a simple MLP-based task-specific head on the latent representation of `[CLS]`, and adopt full-parameter fine-tuning strategy.

## 5. Discussion

In this section, we clarify the significance of Mol-AE through further discussion about four core questions.

### Q1: Encoder-only models are widely used for representation learning in NLP (e.g., BERT). Why don’t they face the inconsistency issues described in Section 3.1?

The main difference lies in the fact that in NLP, words are highly semantic and information-dense; even a single word can convey rich meanings. In contrast, the 3D coordinates in molecules are information-sparse, and the coordinates of an individual atom are meaningless. Therefore, the pretraining objective of reconstructing words in NLP aligns more closely with downstream tasks, as both involve understanding abstract semantics. This alignment is not present in the reconstruction of 3D coordinates.

### Q2: Is the *Identifier* defined in Section 3.2 truly necessary for denoising-based objectives? For instance, CNN-based or GNN-based auto-encoders may not require positional encoding as the *Identifier*

It is essential to emphasize that the *Identifier* defined in Section 3.2 is an abstract concept, representing any information that can identify the object being reconstructed. We use NLP as an example simply because PE in NLP serves as a well-defined *Identifier*. When employing CNN-based models for pixel reconstruction, the *Identifier* is instantiated as the relative spatial relationships between pixels. Similarly, for GNN-based models reconstructing node attributes, the *Identifier* is instantiated as the topological structure that can identify specific nodes.

### Q3: Can we use atom types or 2D graph structure information to play the role of *Identifier* in 3D coordinates?

Indeed, we can use other molecule-related information as the *Identifier*. For instance, we have omitted the modeling of atom types **T** in the preceding text to simplify the formulation, if we consider it, Equation 1 can be rewritten as:

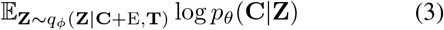

It seems that **T** could serve as the *Identifier* for disrupted coordinates. However, it is crucial to note that this *Identifier* becomes ineffective when two atoms have the same type but are not actually equivalent. Similarly, the strategy of using 2D information as the *Identifier* fails when two atoms are equivalent in 2D structure but not in 3D structure.

Moreover, previous works have not attempted to analyze the challenges of 3D coordinate modeling from the perspective of twisted optimization between *Identifier* and *Content*. Even when attempting to introduce other information alongside 3D modeling, they often overlook this aspect. As a result, a common practice in related work is to perturb 2D, 3D, and atom type information simultaneously (Zhou et al., 2023; Yu et al., 2023), thus failing to satisfy the low-noise requirements of the *Identifier*. The perspective we provide and the simple yet effective approach we adopt can offer new insights into 3D molecular pre-training.

### Q4: What is the relationship between Mol-AE and MAE in vision (He et al., 2022)?

Although Mol-AE employs a similar design to MAE, applying the mask-based paradigm to 3D molecules is not straightforward. Due to significant differences between image and 3D molecular data, current 3D molecular pretraining models are dominated by CD. This has led to two important consensuses: (i) the intensity of disruption should not be too strong, and (ii) biased sequential order information should not be introduced. These factors have made it unlikely for mask-based models similar to MAE to be considered in 3D molecular representation learning. However, we have taken a groundbreaking step by successfully applying the mask-based paradigm to 3D molecular representation learning. Not only have we achieved performance significantly better than CD-based models, but we have also provided compelling evidence to understand why the seemingly contradictory mask-based paradigm works.

## 6. Experiments

### 6.1. Settings

#### Datasets

For pre-training, we use the large-scale molecular dataset provided by Zhou et al. (2023), which contains 19M molecules and 209M conformations generated by ETKGD(Riniker & Landrum, 2015) and Merck Molecular Force Field (Halgren, 1996). Each molecule contains randomly generated 11 conformations in this dataset. For finetuning, we adopt the most widely used benchmark MoleculeNet (Wu et al., 2018), including 9 classification datasets and 6 regression datasets and the data split is the same as Zhou et al. (2023) (cf. Appendix D for more details).

#### Baselines

We use multiple supervised and pre-training methods as our baselines, including supervised and pretraining baselines. D-MPNN (Yang et al., 2019) and AttentiveFP (Xiong et al., 2019) are supervised GNNs methods. N-gram (Liu et al., 2019a), PretrainGNN (Hu et al., 2019), GROVER (Rong et al., 2020), GraphMVP (Liu et al., 2021), MolCLR (Wang et al., 2022b), MoleBLEND (Yu et al., 2023), Uni-Mol (Zhou et al., 2023) are pretraining methods. N-gram embeds the nodes in the graph and assembles them in short walks as the graph representation and Random Forest are used as predictors for downstream tasks. Uni-Mol is the recent SOTA on MoleculeNet benchmark.

#### Implementation Details

We employ Transformer block of hidden size 512, attention heads 64. We set the number of Transformer layers as 15 for encoder, 5 for decoder. For pre-training, We set the drop ratio=0.15 in drop module 𝒟. We use data without hydrogen atoms in pre-training for computational efficiency. We implement positional encoding with sinusoidal PE (Vaswani et al., 2017), and the position of an atom is determined by its order of appearance in SMILES. For downstream evaluation, we only adopt the pre-trained encoder and follow the same fine-tuning protocol of Uni-Mol. For a fair comparison, we evaluate the performance of the official hydrogen-free checkpoint of UniMol, which uses the same pre-training dataset as Mol-AE. (cf. Appendix E for more details about hyper-parameter configuration.)

### 6.2 Main Results

#### Results on Molecular Classification

We present the molecular property classification performance of Mol-AE on 9 widely used tasks. For detailed hyperparameters used in different tasks, please refer to Appendix E. ROC-AUC is employed as the evaluation metric, and the comprehensive results are summarized in Table 1. Mol-AE demonstrates state-of-the-art performance on 6 out of 9 datasets. Moreover, it outperforms Uni-Mol on all tasks. On the largest three datasets, HIV, MUV, and PCBA, Mol-AE exhibits a significant improvement compared to other baselines. Overall, we establish a substantial lead over all other baselines in terms of the average ROC-AUC across all datasets, underscoring the effectiveness of Mol-AE.

**Table 1.** The overall results on 9 molecule classification datasets. We report ROC-AUC score (higher is better) under scaffold splitting. The best results are **bold**. The second-best results are underlined.

| Datasets<br># Molecules | BACE↑<br>1531 | BBBP↑<br>2039 | Tox21↑<br>7831 | SIDER↑<br>1427 | HIV↑<br>41127 | MUV↑<br>93087 | PCBA↑<br>437929 | ClinTox↑<br>1478 | ToxCast↑<br>8575 | Mean↑<br>- |
| --- | --- | --- | --- | --- | --- | --- | --- | --- | --- | --- |
| D-MPNN | 80.9 | 71.0 | 75.9 | 57.0 | 77.1 | 78.6 | 86.2 | 90.6 | 65.5 | 75.87 |
| Attentive FP | 78.4 | 64.3 | 76.1 | 60.6 | 75.7 | 76.6 | 80.1 | 84.7 | 63.7 | 73.36 |
| N-Gram<sub>RF</sub> | 77.9 | 69.7 | 74.3 | <u>66.8</u> | 75.7 | 76.9 | - | 77.5 | - | - |
| PretrainGNN | **84.5** | <u>72.6</u> | 78.1 | 62.7 | <u>79.9</u> | <u>81.3</u> | 86.0 | 72.6 | 65.7 | 75.93 |
| GROVER | 82.6 | 70.0 | 74.3 | 64.8 | 62.5 | 62.5 | 76.5 | 81.2 | 65.4 | 71.09 |
| GraphMVP | 81.2 | 72.4 | 75.9 | 63.9 | 77.0 | 77.7 | - | 79.1 | 63.1 | - |
| MolCLR | 82.4 | 72.2 | 75.0 | 58.9 | 78.1 | 79.6 | - | **91.2** | <u>69.2</u> | - |
| MoleBLEND | 83.7 | **73.0** | 77.8 | 64.9 | 79.0 | 77.2 | - | 87.6 | 66.1 | - |
| Uni-Mol | 83.2 | 71.5 | <u>78.9</u> | 57.7 | 78.6 | 72.6 | <u>88.1</u> | 84.1 | 69.1 | 75.98 |
| MOL-AE | <u>84.1</u> | 72.0 | **80.0** | **67.0** | **80.6** | **81.6** | **88.9** | 87.8 | **69.6** | **79.04** |

#### Results on Molecular Regression

Next, we assess the performance of Mol-AE across 19 molecular regression tasks. Our evaluation employs Mean Absolute Error (MAE) and Root Mean Square Error (RMSE) as the metrics, and the comprehensive results across 6 datasets are presented in Table 2. In cases where datasets include multiple tasks, we compute the mean MAE across all tasks; additional details can be found in Appendix E and D. Mol-AE and achieves the best performance on 5 out of 6 datasets, demonstrating Mol-AE is powerful for molecular regression tasks.

**Table 2.** The overall results on 6 molecule regression datasets. We report Mean Absolute Error (MAE) on QM9, QM8, QM7 tasks and Root Mean Square Error (RMSE) on ESOL, FreeSolv, Lipo tasks under scaffold splitting. The best results are **bold**. The second-best results are underlined. Lower is better on all metrics.

| Datasets<br># Molecules<br># Tasks | QM9↓<br>133885<br>3 | QM8↓<br>21789<br>12 | QM7↓<br>6830<br>1 | ESOL↓<br>1129<br>1 | FreeSolv↓<br>642<br>1 | Lipo↓<br>4200<br>1 |
| --- | --- | --- | --- | --- | --- | --- |
| D-MPNN | 0.0081 | 0.0190 | 103.5 | 1.050 | 2.082 | 0.683 |
| Attentive FP | 0.0081 | 0.0179 | 72.0 | 0.877 | 2.073 | 0.721 |
| N-Gram<sub>RF</sub> | 0.0104 | 0.0236 | 92.8 | 1.074 | 2.688 | 0.812 |
| PretrainGNN | 0.0092 | 0.0200 | 113.2 | 1.100 | 2.764 | 0.739 |
| GROVER | 0.0099 | 0.0218 | 94.5 | 0.983 | 2.176 | 0.817 |
| GraphMVP | - | - | - | 1.029 | - | 0.681 |
| MolCLR | - | 0.0178 | 66.8 | 1.271 | 2.594 | 0.691 |
| MoleBLEND | - | - | - | 0.831 | 1.910 | 0.638 |
| Uni-Mol | <u>0.0054</u> | **0.0160** | <u>58.9</u> | <u>0.844</u> | <u>1.879</u> | <u>0.610</u> |
| MOL-AE | **0.0053** | 0.0161 | **53.8** | **0.830** | **1.448** | **0.607** |

### 6.3 Ablation Study

#### Impact of the Decoder Capacity

First, we explore the effectiveness of using an auto-encoder instead of an encoder-only model. The main difference between these two kinds of model is the capacity of the decoder. Consequently, we investigate how changing the decoder depth affects the performance of Mol-AE. The results are presented in Table 3. We observe a notable decrease in performance when the decoder is too shallow (*L*^dec^≤ 3). This confirms our earlier observations in Section 3, where we noted the inconsistency between pre-training and downstream tasks, with this inconsistency having a smaller impact on middle layers. Additionally, we conduct probing experiments same as Section 3.1 to more intuitively demonstrate that AE outperforms the encoder-only model in molecular understanding tasks, the results are shown in Appendix F.

**Table 3.** Decoder capacity. Using an overly shallow decoder can harm the model’s performance.

| $L^{\text{dec}}$ | Tox21 ↑ | HIV ↑ | QM7 ↓ | FreeSolv ↓ |
| --- | --- | --- | --- | --- |
| 0 | 74.2 | 74.1 | 68.1 | 2.20 |
| 1 | 77.7 | 77.5 | 58.6 | 1.92 |
| 2 | 78.7 | 78.5 | 59.9 | 1.78 |
| 3 | 77.9 | 78.2 | 58.6 | 1.83 |
| 4 | 78.1 | 78.3 | 56.8 | 1.74 |
| 5 | 78.9 | **79.4** | **55.3** | 1.72 |
| 8 | **79.5** | 78.1 | 57.1 | 1.79 |
| 11 | 78.8 | 77.1 | 55.4 | **1.71** |

#### Impact of Order Information Contained in PE

Since PE can not only serve as the *Identifier* but also introduces order information to the model. We investigate how introducing order information might affect the model’s performance. As shown in Table 4, *SMILES* means the atom positions are determined by the order of appearance in the SMILES, while *Random* represents positions determined by a random function. We can find that, compared to the Random approach, utilizing the order from SMILES is more advantageous for modeling the 3D molecular structure. Additionally, the model tends to be more training stable when using PE generated from SMILES (CF. Appendix G for the implementation of random PE and the training process with different PE).

**Table 4.**
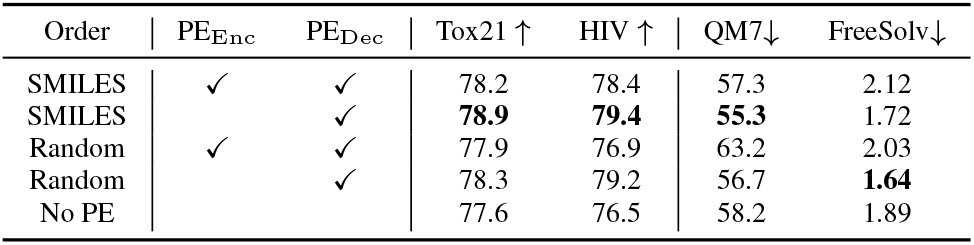
Sequential order information in PE. Introducing PE in encoder will potentially harm the capacity for 3D molecular understanding.

However, directly incorporating such information into the encoder might also introduce biases unrelated to the molecular properties. For example, atoms close in SMILES may not necessarily be close in 3D structure, and a molecule can have multiple valid SMILES. Therefore, we also observe a performance decrease when PE, incorporating order information, is added to the input of the encoder.

#### Impact of Adding PE in Decoder

Firstly, in Table 4, we can observe that introducing PE only in the decoder effectively enhances the performance on downstream tasks. Additionally, in Figure 10 of Appendix H, we illustrate the influence of adding PE on the stability of the training process. We observe that with the addition of PE as an undisturbed *Identifier*, Mol-AE exhibits significantly improved training convergence and stability. We further experiment with introducing PE at intermediate layers within the model to validate the rationale behind solely incorporating PE in the decoder (cf. Appendix I for detailed results).

#### Impact of Different Disruption Methods and Intensity

Based on Mol-AE, we implement a variant named Mol-AE-noise. The only difference between them lies in the strategy used to disrupt input coordinates. Mol-AE employs the dropping strategy, while Mol-AE-noise introduces random noise with the intensity of (0.5^°^*A*, 1^°^*A*, 3^°^*A*, 5^°^*A*). As shown in Table 5, we observe that Mol-AE consistently outperforms Mol-AE-noise. This indicates the effectiveness of using the drop method for disrupting data, allowing the model to focus solely on modeling realistic fragments. We also provide further analysis of influence of different drop ratios in Appendix J, and we find that performance of Mol-AE does not decrease even with high drop ratio (60%).

**Table 5.** Disruption methods. Using dropping to disrupt coordinates could achieve better performance.

| Method | Tox21 ↑ | HIV ↑ | QM7 ↓ | FreeSolv ↓ |
| --- | --- | --- | --- | --- |
| MOL-AE-noise 0.5Å | 78.6 | 79.5 | 56.8 | 1.70 |
| MOL-AE-noise 1Å | 79.5 | 79.9 | 56.6 | 1.68 |
| MOL-AE-noise 3Å | 78.9 | 79.7 | 57.2 | 1.71 |
| MOL-AE-noise 5Å | 78.8 | 79.8 | 56.8 | 1.65 |
| MOL-AE | **80.0** | **80.6** | **53.8** | **1.45** |

## 7. Conclusion

In this paper, we address two common challenges in 3D molecular modeling and provided empirical analyses. To tackle these challenges, we introduced Mol-AE, which leverages an auto-encoder to mitigate potential inconsistencies between pre-training and downstream tasks. Additionally, by carefully discussing the properties of the *Content* and *Identifier* roles, we proposed a new objective, the 3D Cloze Test, to train the model for better molecular understanding. Extensive experiments demonstrated the superior performance of Mol-AE in 3D molecular understanding.

## Acknowledgements

We would like to thank Qiying Yu from AIR and Zequn Liu from PKU for their insightful discussions on the project. We also thank other members from AIR for their valuable feedback given during the internal seminar. This work is supported by the National Science and Technology Major Project (2022ZD0117502), the National Natural Science Foundation of China (62276002 and 62376133), and the PharMolix Inc.

## Impact Statement

Our work can help the AI4Science field better understand and develop robust molecular representation learning models. With the increasing application of molecular representation learning models in various scenarios, designing a more powerful molecular representation learning model has become a crucial aspect driving progress in the field. This work reveals the shortcomings of existing models through analysis and provides insights for designing better molecular representation learning models, which holds significant practical implications. However, we also acknowledge that this work inherits the potential negative impacts of existing molecular pre-training models, such as the possibility of being used to design and manufacture molecules with biological hazards.

## A. Related Work

Early approaches to molecular representation learning predominantly focused on 1D SMILES (Wang et al., 2019; Chithrananda et al., 2020; Guo et al., 2021; Honda et al., 2019) and 2D graphs (Li et al., 2021; Lu et al., 2021; Fang et al., 2022b; Xia et al., 2022). Recently, there has been a growing interest in 3D molecular data, which could provide a more comprehensive reflection of physical properties, including information not captured by 1D and 2D data, such as conformation details.

Recent developments in 3D modeling involve self-supervised learning directly from unlabeled 3D data to learn informative features (Liu et al., 2022a; Stärk et al., 2022; Zhou et al., 2023; Yu et al., 2023; Feng et al., 2023).

Regarding model structure, most 3D molecular representation learning has used encoder-only methods, which include Transformer-based encoders and GNN-based encoders. For Transformer-based models, a common approach is to encode the relative positions of atoms as attention bias to enable the model to understand 3D information (Zhou et al., 2023; Yu et al., 2023; Luo et al., 2022). For GNN-based encoder models, a prevalent method involves treating relative atom information as edge features and utilizing message passing (Gilmer et al., 2017) for representation learning (Feng et al., 2023).

Regarding pre-training objectives, the primary methods include geometry prediction and coordinate denoising. In the case of geometry prediction, models are trained to predict intrinsic geometric properties of molecules, such as bond lengths, bond angles (Fang et al., 2022a), shortest paths (Luo et al., 2022; Yu et al., 2023), node types (Zhou et al., 2023), and more. For coordinate denoising, the approach involves introducing random noise to the input coordinates, and then training the model to denoise them using an SE(3) head to recover the original coordinates (Luo et al., 2022; Yu et al., 2023). Additionally, coordinate denoising is often combined with distance reconstruction (Zhou et al., 2023) to achieve enhanced model performance.

## B. Performance of Uni-Mol with PE on Downstream Tasks

We modify the original Uni-Mol model by adding positional encoding to the representation of each atom to assist the model in better distinguishing between different atoms. And this approach is referred to as Uni-Mol-PE. As shown in Figure 5(a), compared to the original Uni-Mol model, the Uni-Mol-PE model exhibits lower reconstruction errors, smaller loss fluctuations, and better convergence during pre-training. This indicates that introducing positional encoding can indeed help the model distinguish between different atoms to reconstruct the corrupted structural information and compensate for the disrupted original identifier.

**Figure 5.**
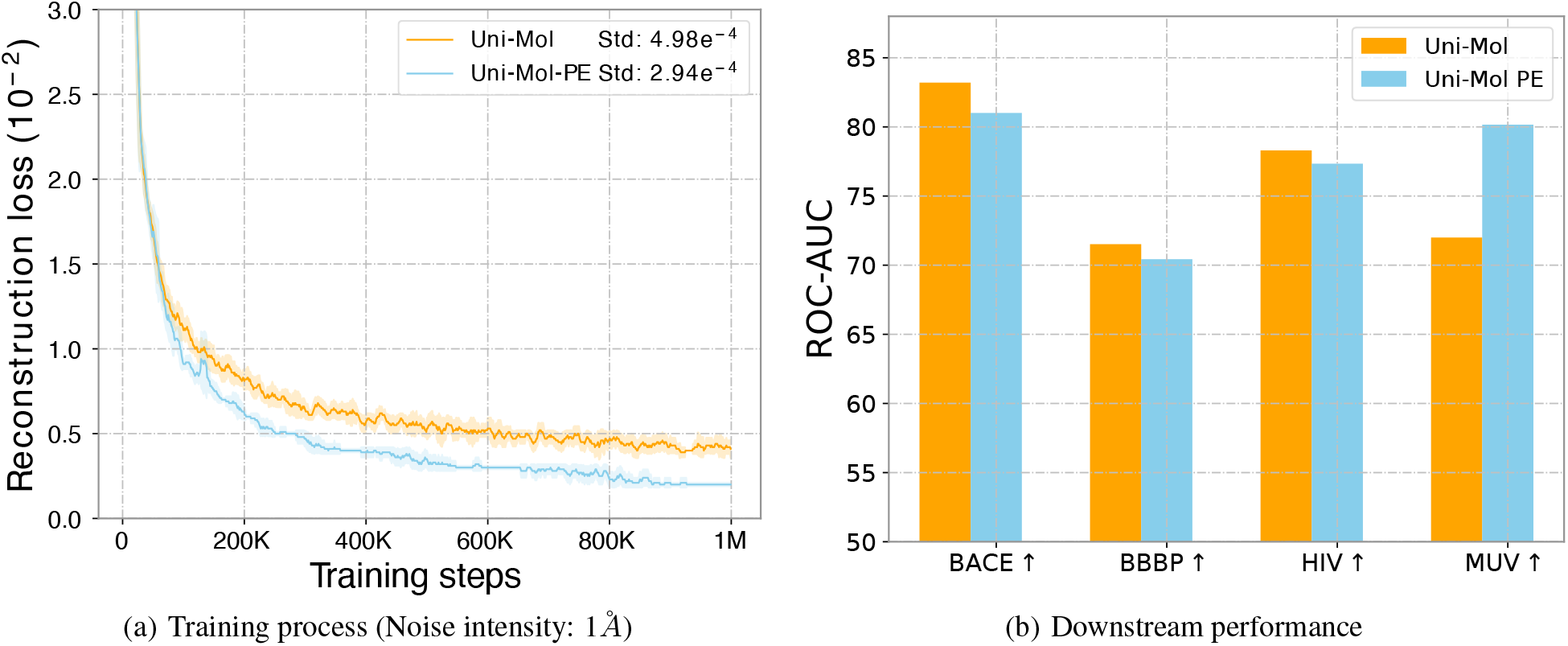
Impact of introducing PE as additional Identifier to vanilla Uni-Mol.

However, when comparing the performance of the Uni-Mol model and the Uni-Mol-PE model on different downstream tasks (shown in Figure 5(b)), we find that the Uni-Mol-PE model exhibits a decrease in performance compared to the Uni-Mol model on several downstream tasks. This indicates that directly incorporating the atomic order information contained in the SMILES into the model may not always be advantageous for the model in molecular representation learning. This is because the atomic order information contained in the SMILES may be biased, and providing a predefined order for different atoms, similar to NLP models, may not necessarily be helpful in learning a good molecular representation.

## C. Analysis of Transformer-M

We perform the same analytic experiments as Section 3 on another widely used *EnCD* 3D molecular pre-training model, Transformer-M (Luo et al., 2022). The observed phenomena closely resemble those of Uni-Mol. The results are shown in Figure 6.

**Figure 6.**
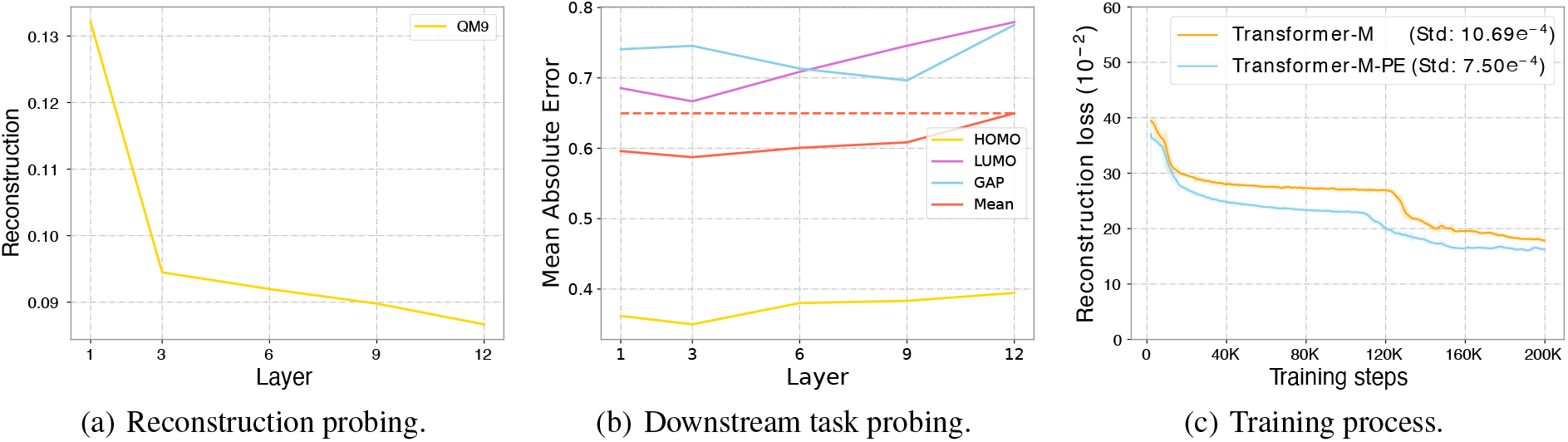
Transformer-M also faces inconsistency problem and twisted optimization problem.

- **Transformer-M also struggles with the impact of inconsistency (same as Uni-Mol in Section 3.1)**. Since Transformer-M only provides source codes for fine-tuning on QM9 dataset, we conduct reconstruction probing on QM9 (Figure 6.a) and downstream performance probing on three representative downstream tasks of QM9 (Figure 6.b, the metric is Mean Absolute Error, lower is better). The results indicate that deeper representations in Transformer-M excel at reconstruction but exhibit progressively poorer downstream performance, indicating the same inconsistency problem within Transformer-M. Therefore, directly fine-tuning the entire model on downstream tasks with an encoder-only model may affect performance.
- **Transformer-M also faces the issue of twisted optimization (same as Uni-Mol in Section 3.2)**. Similar to Uni-Mol, Transformer-M also treats the noisy atomic structural information as identifiers for different atoms, leading to difficulty in distinguishing different atoms after noise addition, thus causing higher reconstruction loss. To address this issue, we introduce sequential position encoding determined by the order of atoms in SMILES for the Transformer-M model, referred to as Transformer-M-PE. The comparison reveals that Transformer-M-PE exhibits significantly lower reconstruction loss compared to the Transformer-M model (Figure 6.c).

It’s worth noting that Transformer-M utilizes a completely different pre-training dataset and fine-tuning protocol from Uni-Mol, thus can strongly confirm the universality of the observed two phenomena in EnCD models.

## D. Datasets

### Pre-training Datasets

We use the dataset provided by Zhou et al. (2023), which contains 19M molecules and 209M conformations generated by ETKGD (Riniker & Landrum, 2015) and Merck Molecular Force Field (Halgren, 1996). During the pre-training process, to ensure training efficiency, we remove all hydrogen atoms in the pre-training dataset.

### Fine-tuning Datasets

We conduct experiments on the MoleculeNet(Wu et al., 2018) benchmark in the molecular property prediction task. MoleculeNet serves as a widely recognized benchmark in the field of molecular property prediction. Here, we offer the statistics and basic information of the MoleculeNet benchmark datasets in Table 6.

**Table 6.**
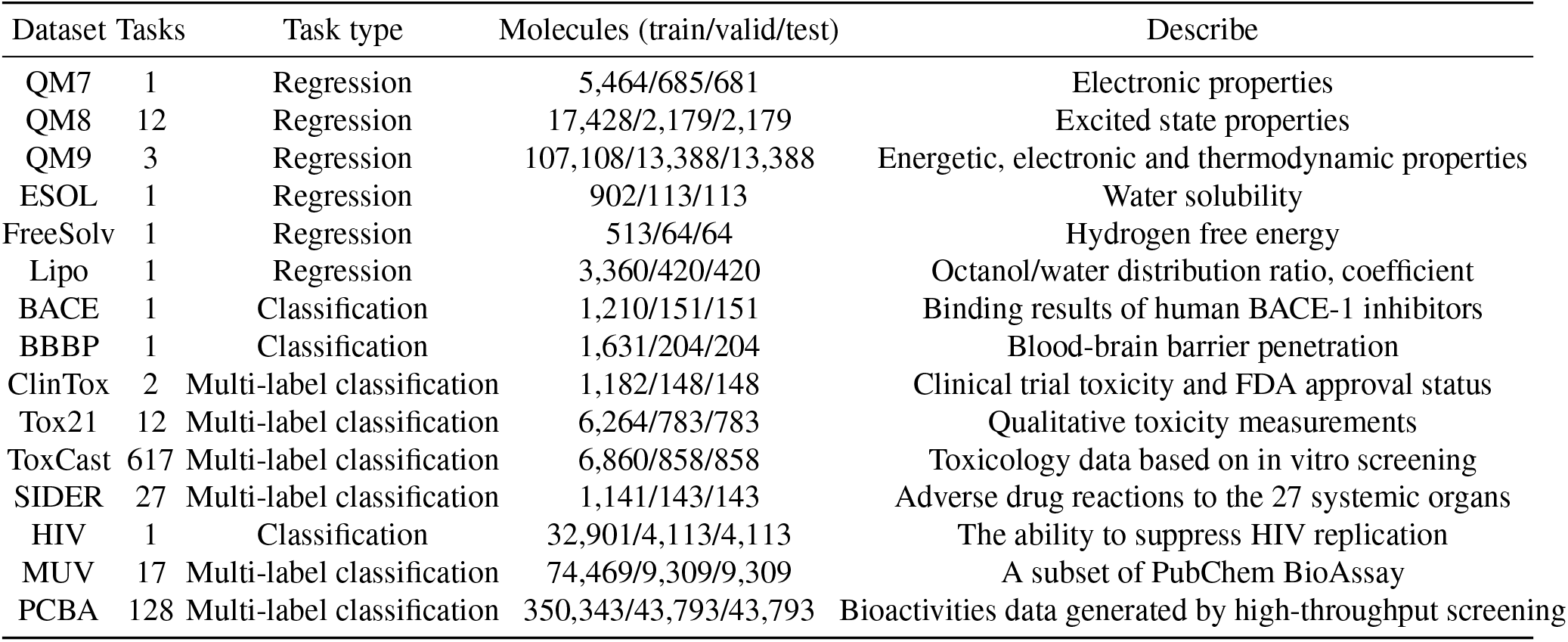
Summary information of the MoleculeNet benchmark datasets.

| Dataset | Tasks | Task type | Molecules (train/valid/test) | Describe |
| --- | --- | --- | --- | --- |
| QM7 | 1 | Regression | 5,464/685/681 | Electronic properties |
| QM8 | 12 | Regression | 17,428/2,179/2,179 | Excited state properties |
| QM9 | 3 | Regression | 107,108/13,388/13,388 | Energetic, electronic and thermodynamic properties |
| ESOL | 1 | Regression | 902/113/113 | Water solubility |
| FreeSolv | 1 | Regression | 513/64/64 | Hydrogen free energy |
| Lipo | 1 | Regression | 3,360/420/420 | Octanol/water distribution ratio, coefficient |
| BACE | 1 | Classification | 1,210/151/151 | Binding results of human BACE-1 inhibitors |
| BBBP | 1 | Classification | 1,631/204/204 | Blood-brain barrier penetration |
| ClinTox | 2 | Multi-label classification | 1,182/148/148 | Clinical trial toxicity and FDA approval status |
| Tox21 | 12 | Multi-label classification | 6,264/783/783 | Qualitative toxicity measurements |
| ToxCast | 617 | Multi-label classification | 6,860/858/858 | Toxicology data based on in vitro screening |
| SIDER | 27 | Multi-label classification | 1,141/143/143 | Adverse drug reactions to the 27 systemic organs |
| HIV | 1 | Classification | 32,901/4,113/4,113 | The ability to suppress HIV replication |
| MUV | 17 | Multi-label classification | 74,469/9,309/9,309 | A subset of PubChem BioAssay |
| PCBA | 128 | Multi-label classification | 350,343/43,793/43,793 | Bioactivities data generated by high-throughput screening |

### Evaluation protocol of QM9

QM9 contains several quantum mechanical properties of different quantitative ranges, and we select *homo, lumo and gap* of similar quantitative range, following the setup of the previous work (Zhou et al., 2023).

## E. Hyper-Parameter Configuration

We implement Mol-AE using 15 stacked Transformer layers in encoder and 5 stacked Transformer layers in decoder, each with 64 attention heads. The model dimension and feedforward dimension of each Transformer layer are 512 and 2048. The total number of Mol-AE’s parameters is 48M. We use Adam (Kingma & Ba, 2014) and polynomial learning rate scheduler to train Mol-AE and set the learning rate 1e-4, weight decay 1e-4, warmup step 10K. The total training step is 1M and each batch has 128 samples at maximum. We train Mol-AE on a single NVIDIA A100 GPU for about 2 days.

For more pre-training hyper-parameters, please refer to Table 7.

**Table 7.** Mol-AE hyper-parameters for pre-training.

| Hyper-parameters | Value |
| --- | --- |
| Learning rate | 1e-4 |
| LR scheduler | polynomial_decay |
| Warmup updates | 10K |
| Max updates | 1M |
| Batch size | 128 |
| Distance loss function and its weight | Smooth L1, 10.0 |
| Coordinate loss function and its weight | Smooth L1, 5.0 |
| Atom loss function and its weight | Cross entropy, 1.0 |
| FFN dropout | 0.1 |
| Attention dropout | 0.1 |
| Embedding dropout | 0.1 |
| Num of encoder layers | 15 |
| Num of encoder attention heads | 64 |
| Encoder embedding dim | 512 |
| Encoder FFN dim | 2048 |
| Num of decoder layers | 5 |
| Num of decoder attention heads | 64 |
| Decoder embedding dim | 512 |
| Decoder FFN dim | 2048 |
| Adam ($\beta_1, \beta_2$) | (0.9, 0.99) |
| Drop ratio | 0.15 |
| Vocabulary size (atom types) | 30 |
| Activation function | GELU |

In different downstream task, we use different hyper-parameters. For detailed fine-tuning hyper-parameters, please refer to Table 8.

**Table 8.** Mol-AE hyper-parameters for fine-tuning.

| Tasks | Epochs | Batch size | Learning rate | Warmup Ratio | Dropout | Pooler-dropout |
| --- | --- | --- | --- | --- | --- | --- |
| BACE | 20 | 64 | 1e-4 | 0.36 | 0.1 | 0.2 |
| BBBP | 40 | 128 | 4e-4 | 0.18 | 0.1 | 0.1 |
| TOX21 | 80 | 128 | 1e-4 | 0.06 | 0.1 | 0.1 |
| SIDER | 40 | 32 | 5e-4 | 0.5 | 0.1 | 0 |
| HIV | 5 | 256 | 5e-5 | 0.1 | 0.1 | 0.2 |
| MUV | 20 | 128 | 2e-5 | 0.3 | 0.1 | 0.1 |
| PCBA | 20 | 128 | 1e-4 | 0.06 | 0.1 | 0.1 |
| ClinTox | 80 | 256 | 5e-5 | 0.25 | 0.1 | 0.7 |
| ToxCast | 160 | 64 | 1e-4 | 0.06 | 0.1 | 0.2 |
| QM9 | 40 | 128 | 1e-4 | 0.06 | 0.1 | 0 |
| QM8 | 120 | 32 | 1e-4 | 0.02 | 0 | 0 |
| QM7 | 200 | 32 | 3e-4 | 0.06 | 0.1 | 0.1 |
| ESOL | 200 | 256 | 5e-4 | 0.06 | 0.1 | 0.4 |
| FreeSolv | 160 | 64 | 8e-5 | 0.1 | 0.1 | 0.4 |
| Lipo | 100 | 64 | 1e-4 | 0.24 | 0.1 | 0.1 |

## F. More Strict Validation of The Capability of Auto-Encoder

Here, we conduct more straightforward experiments to better demonstrate that using the AE model can effectively escape from the negative impacts caused by inconsistency problem. Specifically, we carry out the two probing experiments and one fine-tuning experiment on Mol-AE and Mol-AE-noise. Mol-AE-noise is a small variant of Mol-AE. The only difference between them lies in the strategy used to disrupt input coordinates. Mol-AE employs the dropping strategy, while Mol-AE-noise introduces random noise. Both Mol-AE and Mol-AE-noise contain 15 encoder layers and 5 decoder layers.

### F.1. Probing Experiments on Mol-AE

Similar to layer-wise probing in section 3.1, we conduct the same two probing experiments on Mol-AE (fix the whole model and only finetune task head). Due to the absence of representations for dropped atoms in Mol-AE’s encoder, we probe representations solely from layer 16 to layer 20. The reconstruction loss is shown in Figure 7(a), and the downstream performance is detailed in Figure 7(b). It’s shown that in Mol-AE, as the layer depth increases, the corresponding representations performs better in reconstructing coordinates (pre-training task) but worse in downstream tasks. This underscores the necessity of adopting the AE structure and omitting the decoder in downstream tasks.

**Figure 7.**
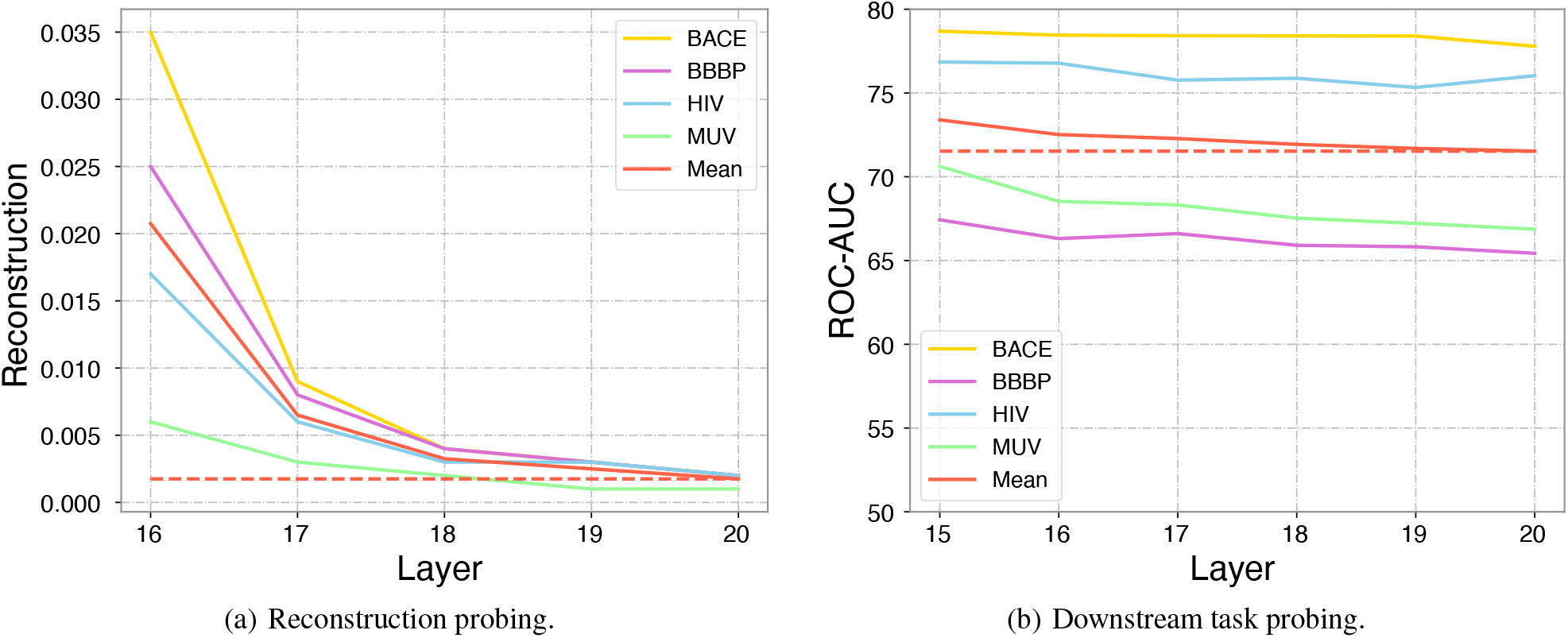
Probing experiments on Mol-AE.

### F.2. Probing Experiments on Mol-AE-noise

In Mol-AE-noise, since all atoms in the encoder have valid representations, we can conduct the same layer-wise probing on representations from layers 1-20. The reconstruction loss is presented in Figure 8(a), and the downstream performance is detailed in Figure 8(b). The results indicate that since Mol-AE-noise introduces *Identifier* in the decoder, there is a clearer division of labor between its encoder and decoder. Specifically, compared to Uni-Mol (Figure 2), the reconstruction loss of the last five layers in Mol-AE-noise decreases more rapidly and the generalization to downstream tasks of each encoder layer in the Mol-AE-noise is more stable. This indicates that pre-training with an AE structure and abandoning the decoder in downstream tasks is more advantageous for the model to escape from the negative impacts caused by inconsistency.

**Figure 8.**
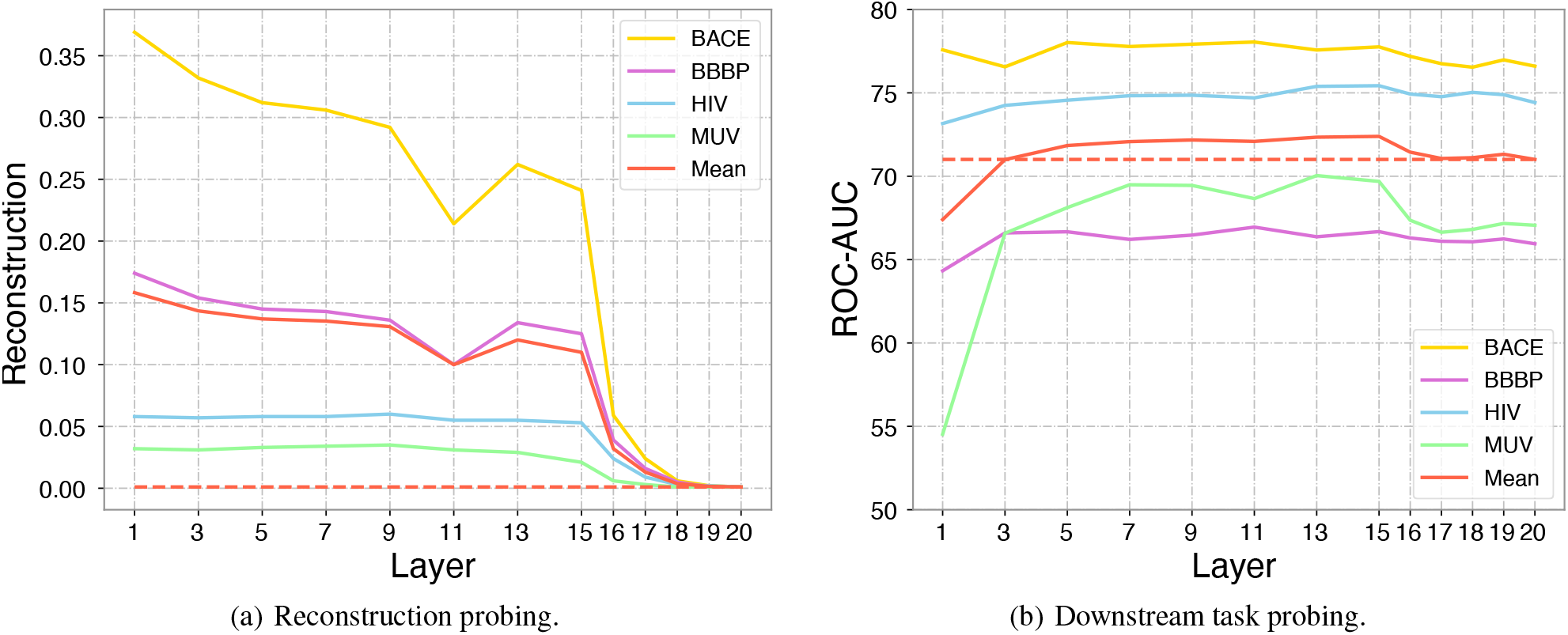
Probing experiments on Mol-AE-noise.

### F.3. Fine-tuning Experiments on Mol-AE

We also examine the impact of fine-tuning only the encoder of Mol-AE versus fine-tuning both the encoder and decoder of Mol-AE (Mol-AE-full) on downstream task, as shown in Table 9. It can be observed that despite Mol-AE-full having more learnable parameters, its performance consistently lags behind Mol-AE on downstream tasks, indicating the necessity of using an AE model for pre-training and removing the decoder in downstream tasks.

**Table 9.**
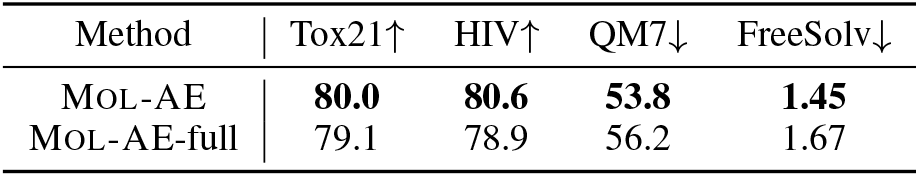
Performance comparison of Mol-AE and Mol-AE-full on downstream tasks.

## G. Order Information Contained in PE

To generate a PE determined by a random function, we first choose a threshold, *max len*, to ensure that the number of atoms in each molecule does not exceed *max len*. Then, before the model training begins, we use *np*.*random*.*permutation(max len)* to instantiate a random mapping function *f*_idx_ : [1, max_len] ↦ [1, max_len]. This random mapping is fixed once instantiated. During training, if an atom has a position *i* in the SMILES, then its random PE is PE(*f*_idx_(*i*)). As shown in **Figure 9**, when using Random PE, the model training exhibits larger fluctuations, and the final reconstruction loss is higher.

**Figure 9.**
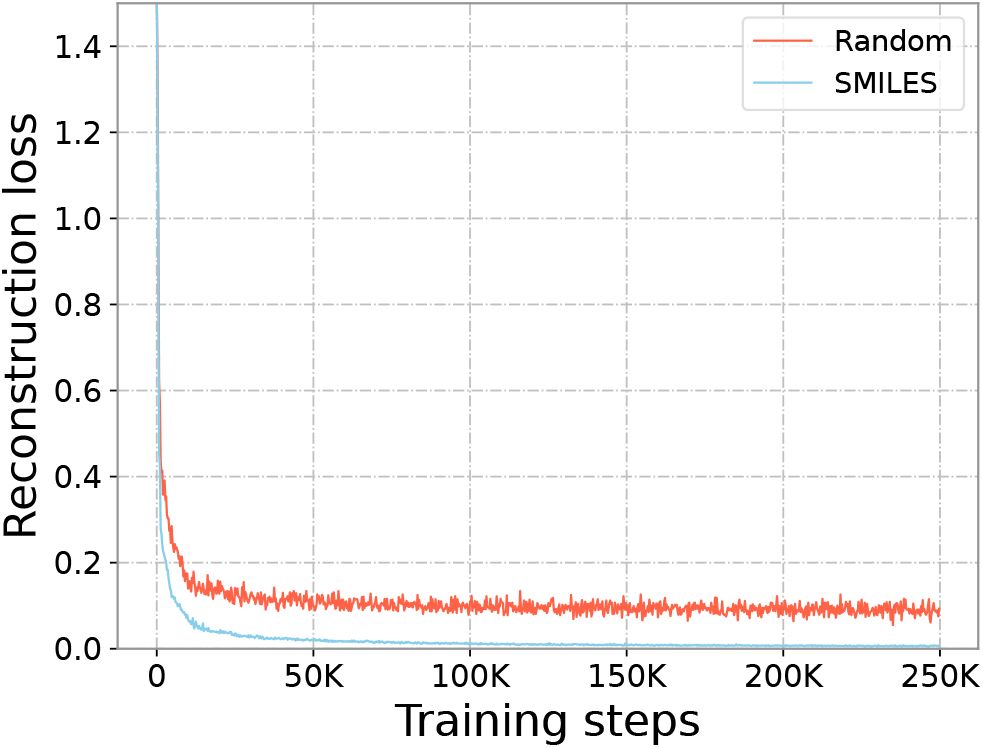
Training process of Mol-AE when different PE is adopted.

**Figure 10.**
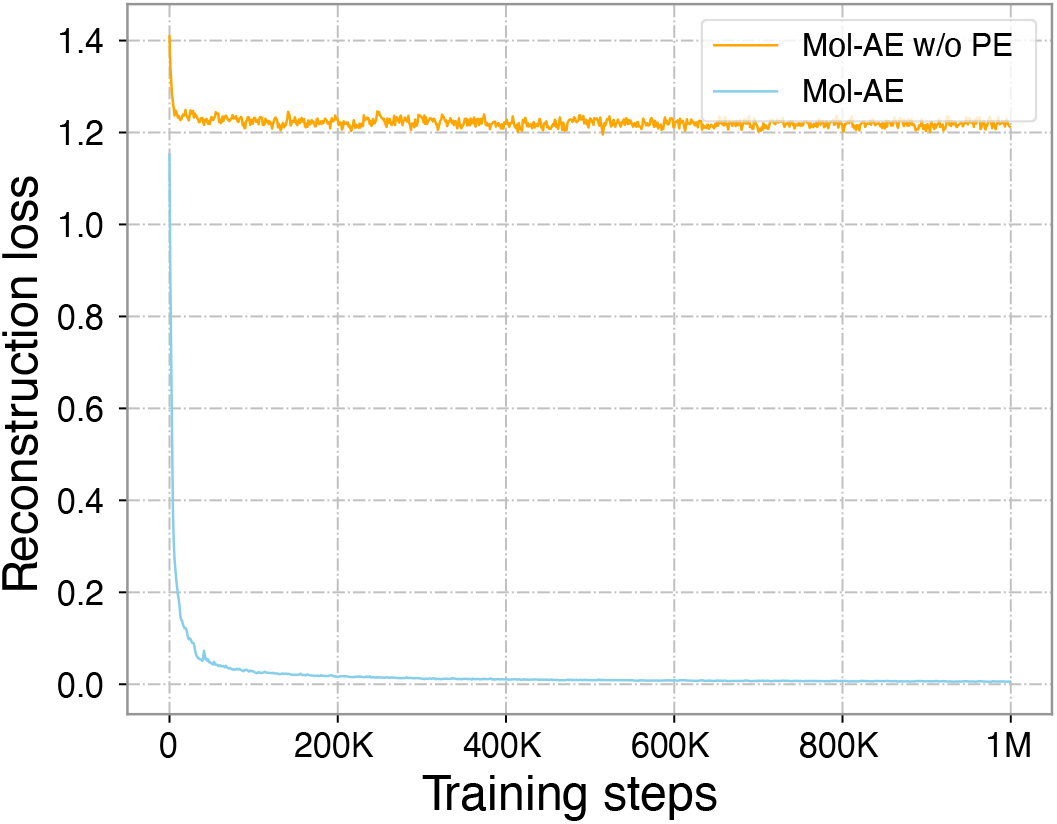
When PE is added to Mol-AE, the training convergence and stability are significantly improved.

## H. Impact of Adding PE in Decoder

We train a new Mol-AE model without PE in decoder (Mol-AE w/o PE) and compare the training curve of Mol-AE w/o PE and that of Mol-AE. We can find that the addition of PE in decoder will significantly improve the reconstruction ability of the model and further improve the convergence in the pre-training process, as shown in Figure 10.

## I. Ablation Study on Adding PE to Different Layers

We conduct an ablation study to evaluate which layer benefits more from the addition of PE. Specifically, in the AE model consisting of 15 encoder layers and 5 decoder layers, we individually attempt to add PE to the outputs of different layers of the model (where layer=0 indicates that PE is added to the input of the entire model) and report the performance of fine-tuning encoder on downstream tasks. The results are presented in Table 10. We observe two interesting phenomena:

**Table 10.** Ablation study on adding PE to different layers.

| Data | Layer 0 | Layer 5 | Layer 10 | Layer 15 | Layer 16 | Layer 17 | Layer 18 | Layer 19 | Layer 20 |
| --- | --- | --- | --- | --- | --- | --- | --- | --- | --- |
| Tox21 ↑ | 78.2 | 77.9 | 77.4 | **78.9** | 78.6 | 78.9 | 77.6 | 77.1 | 77.3 |
| HIV ↑ | 78.4 | 78.1 | 77.6 | <u>79.4</u> | 79.3 | **79.7** | 78.6 | 79.1 | 78.3 |
| QM7 ↓ | 57.2 | 58.1 | 59.4 | **55.3** | <u>55.4</u> | 56.9 | 57.7 | 57.4 | 57.8 |
| FreeSolve ↓ | 2.11 | 2.13 | 2.15 | <u>1.72</u> | **1.69** | 1.77 | 1.73 | 1.76 | 1.84 |

- If PE is incorporated into the encoder, the performance consistently becomes worse in downstream tasks. Moreover, the closer PE is incorporated to the latent representations, the more pronounced performance degradation. This could be attributed to the sequential order bias contained in PE would have a greater negative impact when PE is too close to the latent representations in encoder.
- If PE is added to the decoder, there is no significant change in downstream performance when the layer where PE is added is relatively close to the latent representations (e.g., Layer 16, 17).

However, for simplification, when we attempt to add PE in the decoder, we directly incorporate PE into the latent representations, as many Seq2Seq models typically incorporate PE directly into the decoder’s input (Raffel et al., 2020; Lewis et al., 2019), which is similar to our approach of adding PE to the latent representation.

## J. Impact of Drop Ratio

Here, we evaluate how different drop ratio would affect the model’s performance. The results are show in Table 11. We can observe that even with a drop ratio of 60% (which is really high for molecules), the performance of Mol-AE does not obviously decrease (still better than Uni-Mol). However, when drop ratio=0.6, under the Transformer architecture, the floating-point operations performed is approximately only 22% compared to when the dropratio=0.15 (0.4^2^*/*0.85^2^). Such acceleration suggests that Mol-AE may hold great potential for large molecule modeling.

**Table 11.** The impact of drop ratio on downstream performance.

| Dataset | Drop 7% | Drop 15% | Drop 30% | Drop 45% | Drop 60% | Drop 75% | Drop 90% | Uni-Mol |
| --- | --- | --- | --- | --- | --- | --- | --- | --- |
| Tox21↑ | 79.2 | **80.0** | <u>80.0</u> | 79.4 | 79.2 | 78.2 | 75.3 | 78.9 |
| HIV↑ | 79.6 | **80.6** | <u>79.1</u> | 79.0 | <u>80.1</u> | 78.3 | 76.3 | 78.6 |
| QM7↓ | <u>53.6</u> | 53.8 | **52.8** | 55.3 | <u>56.0</u> | 60.7 | 67.2 | 58.9 |
| FreeSolv↓ | 1.49 | **1.45** | <u>1.47</u> | 1.61 | 1.66 | 2.02 | 2.31 | 1.88 |

## K. Details of SE(3)-equivariant head

The SE(3)-equivariant head in Mol-AE refers to the coordinate prediction head that is equivariant under transformations in SE(3) group, such as 3D translations and rotations, which are essential for 3D spatial tasks. We use the same SE(3)-equivariant head from Uni-Mol (Zhou et al., 2023). The head can be formulated as follows:

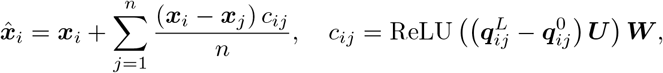

Here, ***x***_*i*_ is the *i*-th atom’s coordinates in the input molecule, and 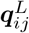 is the Pair-Wise Feature between *i*-th atom and *j*-th atom at the *L*-th layer. ***U*** ∈ ℝ^*H*×*H*^ and ***W*** ∈ ℝ^*H*×1^ are the projection matrices.

## Footnotes

1 The formula can be derived from the standard ELBO by omitting the KL term.

2 For example, in SMILES C=O, the oxygen atom is the second atom, thus its position is 2.

## References

Adelusi, T. I., Oyedele, A.-Q. K., Boyenle, I. D., Ogunlana, A. T., Adeyemi, R. O., Ukachi, C. D., Idris, M. O., Olaoba, O. T., Adedotun, I. O., Kolawole, O. E., et al. Molecular modeling in drug discovery. Informatics in Medicine Unlocked, 29:100880, 2022.

Ba, J. L., Kiros, J. R., and Hinton, G. E. Layer normalization. arXiv preprint arXiv:1607.06450, 2016.

Cai, X., Huang, J., Bian, Y., and Church, K. Isotropy in the contextual embedding space: Clusters and manifolds. In International Conference on Learning Representations, 2020.

Chithrananda, S., Grand, G., and Ramsundar, B. Chemberta: large-scale self-supervised pretraining for molecular property prediction. arXiv preprint arXiv:2010.09885, 2020.

Devlin, J., Chang, M.-W., Lee, K., and Toutanova, K. Bert: Pre-training of deep bidirectional transformers for language understanding. arXiv preprint arXiv:1810.04805, 2018.

Ethayarajh, K. How contextual are contextualized word representations? comparing the geometry of bert, elmo, and gpt-2 embeddings. arXiv preprint arXiv:1909.00512, 2019.

Fang, X., Liu, L., Lei, J., He, D., Zhang, S., Zhou, J., Wang, F., Wu, H., and Wang, H. Geometry-enhanced molecular representation learning for property prediction. Nature Machine Intelligence, 4(2):127–134, 2022a.

Fang, Y., Zhang, Q., Yang, H., Zhuang, X., Deng, S., Zhang, W., Qin, M., Chen, Z., Fan, X., and Chen, H. Molecular contrastive learning with chemical element knowledge graph. In Proceedings of the AAAI Conference on Artificial Intelligence, volume 36, pp. 3968–3976, 2022b.

Feng, S., Ni, Y., Lan, Y., Ma, Z.-M., and Ma, W.-Y. Fractional denoising for 3d molecular pre-training. In International Conference on Machine Learning, pp. 9938–9961. PMLR, 2023.

Gastegger, M., Schütt, K. T., and Müller, K.-R. Machine learning of solvent effects on molecular spectra and reactions. Chemical science, 12(34):11473–11483, 2021.

Gilmer, J., Schoenholz, S. S., Riley, P. F., Vinyals, O., and Dahl, G. E. Neural message passing for quantum chemistry. In International conference on machine learning, pp. 1263–1272. PMLR, 2017.

Guo, Z., Sharma, P., Martinez, A., Du, L., and Abraham, R. Multilingual molecular representation learning via contrastive pre-training. arXiv preprint arXiv:2109.08830, 2021.

Halgren, T. A. Merck molecular force field. i. basis, form, scope, parameterization, and performance of mmff94. Journal of computational chemistry, 17(5-6):490–519, 1996.

He, K., Chen, X., Xie, S., Li, Y., Dollár, P., and Girshick, R. Masked autoencoders are scalable vision learners. In Proceedings of the IEEE/CVF conference on computer vision and pattern recognition, pp. 16000–16009, 2022.

Honda, S., Shi, S., and Ueda, H. R. Smiles transformer: Pre-trained molecular fingerprint for low data drug discovery. arXiv preprint arXiv:1911.04738, 2019.

Hu, W., Liu, B., Gomes, J., Zitnik, M., Liang, P., Pande, V., and Leskovec, J. Strategies for pre-training graph neural networks. arXiv preprint arXiv:1905.12265, 2019.

Jiao, R., Han, J., Huang, W., Rong, Y., and Liu, Y. Energy-motivated equivariant pretraining for 3d molecular graphs. In Proceedings of the AAAI Conference on Artificial Intelligence, volume 37, pp. 8096–8104, 2023.

Ju, W., Liu, Z., Qin, Y., Feng, B., Wang, C., Guo, Z., Luo, X., and Zhang, M. Few-shot molecular property prediction via hierarchically structured learning on relation graphs. Neural Networks, 163:122–131, 2023.

Kingma, D. P. and Ba, J. Adam: A method for stochastic optimization. arXiv preprint arXiv:1412.6980, 2014.

Lewis, M., Liu, Y., Goyal, N., Ghazvininejad, M., Mohamed, A., Levy, O., Stoyanov, V., and Zettlemoyer, L. Bart: Denoising sequence-to-sequence pre-training for natural language generation, translation, and comprehension. arXiv preprint arXiv:1910.13461, 2019.

Li, P., Wang, J., Qiao, Y., Chen, H., Yu, Y., Yao, X., Gao, P., Xie, G., and Song, S. An effective self-supervised framework for learning expressive molecular global representations to drug discovery. Briefings in Bioinformatics, 22 (6):bbab109, 2021.

Liu, S., Demirel, M. F., and Liang, Y. N-gram graph: Simple unsupervised representation for graphs, with applications to molecules. Advances in neural information processing systems, 32, 2019.

Liu, S., Wang, H., Liu, W., Lasenby, J., Guo, H., and Tang, J. Pre-training molecular graph representation with 3d geometry. arXiv preprint arXiv:2110.07728, 2021.

Liu, S., Guo, H., and Tang, J. Molecular geometry pretraining with se (3)-invariant denoising distance matching. arXiv preprint arXiv:2206.13602, 2022a.

Liu, Y., Ott, M., Goyal, N., Du, J., Joshi, M., Chen, D., Levy, O., Lewis, M., Zettlemoyer, L., and Stoyanov, V. Roberta: A robustly optimized BERT pretraining approach. CoRR, abs/1907.11692, 2019b. URL http://arxiv.org/abs/1907.11692.

Liu, Y., Wang, L., Liu, M., Lin, Y., Zhang, X., Oztekin, B., and Ji, S. Spherical message passing for 3d molecular graphs. In International Conference on Learning Representations (ICLR), 2022b.

Lu, Y., Jiang, X., Fang, Y., and Shi, C. Learning to pretrain graph neural networks. In Proceedings of the AAAI conference on artificial intelligence, volume 35, pp. 4276–4284, 2021.

Luo, S., Chen, T., Xu, Y., Zheng, S., Liu, T.-Y., Wang, L., and He, D. One transformer can understand both 2d & 3d molecular data. arXiv preprint arXiv:2210.01765, 2022.

Pinzi, L. and Rastelli, G. Molecular docking: shifting paradigms in drug discovery. International journal of molecular sciences, 20(18):4331, 2019.

Raffel, C., Shazeer, N., Roberts, A., Lee, K., Narang, S., Matena, M., Zhou, Y., Li, W., and Liu, P. J. Exploring the limits of transfer learning with a unified text-to-text transformer. Journal of machine learning research, 21 (140):1–67, 2020.

Riniker, S. and Landrum, G. A. Better informed distance geometry: using what we know to improve conformation generation. Journal of chemical information and modeling, 55(12):2562–2574, 2015.

Rong, Y., Bian, Y., Xu, T., Xie, W., Wei, Y., Huang, W., and Huang, J. Self-supervised graph transformer on large-scale molecular data. Advances in Neural Information Processing Systems, 33:12559–12571, 2020.

Satorras, V. G., Hoogeboom, E., and Welling, M. E (n) equivariant graph neural networks. In International conference on machine learning, pp. 9323–9332. PMLR, 2021.

Scholkopf, B., Sung, K.-K., Burges, C. J., Girosi, F., Niyogi, P., Poggio, T., and Vapnik, V. Comparing support vector machines with gaussian kernels to radial basis function classifiers. IEEE transactions on Signal Processing, 45 (11):2758–2765, 1997.

Schwaller, P., Vaucher, A. C., Laino, T., and Reymond, J.-L. Prediction of chemical reaction yields using deep learning. Machine learning: science and technology, 2 (1):015016, 2021.

Stärk, H., Beaini, D., Corso, G., Tossou, P., Dallago, C., Günnemann, S., and Liò, P. 3d infomax improves gnns for molecular property prediction. In International Conference on Machine Learning, pp. 20479–20502. PMLR, 2022.

Tenney, I., Xia, P., Chen, B., Wang, A., Poliak, A., McCoy, R. T., Kim, N., Van Durme, B., Bowman, S. R., Das, D., et al. What do you learn from context? probing for sentence structure in contextualized word representations. arXiv preprint arXiv:1905.06316, 2019.

van Tilborg, D., Alenicheva, A., and Grisoni, F. Exposing the limitations of molecular machine learning with activity cliffs. Journal of Chemical Information and Modeling, 62(23):5938–5951, 2022.

Vaswani, A., Shazeer, N., Parmar, N., Uszkoreit, J., Jones, L., Gomez, A. N., Kaiser, Ł., and Polosukhin, I. Attention is all you need. Advances in neural information processing systems, 30, 2017.

Wang, L., Zhou, Y., Wang, Y., Zheng, X., Huang, X., and Zhou, H. Regularized molecular conformation fields. Advances in Neural Information Processing Systems, 35: 18929–18941, 2022a.

Wang, S., Guo, Y., Wang, Y., Sun, H., and Huang, J. Smilesbert: large scale unsupervised pre-training for molecular property prediction. In Proceedings of the 10th ACM international conference on bioinformatics, computational biology and health informatics, pp. 429–436, 2019.

Wang, Y., Wang, J., Cao, Z., and Barati Farimani, A. Molecular contrastive learning of representations via graph neural networks. Nature Machine Intelligence, 4(3):279–287, 2022b.

Wu, Z., Ramsundar, B., Feinberg, E. N., Gomes, J., Geniesse, C., Pappu, A. S., Leswing, K., and Pande, V. Moleculenet: a benchmark for molecular machine learning. Chemical science, 9(2):513–530, 2018.

Xia, J., Zhao, C., Hu, B., Gao, Z., Tan, C., Liu, Y., Li, S., and Li, S. Z. Mole-bert: Rethinking pre-training graph neural networks for molecules. In The Eleventh International Conference on Learning Representations, 2022.

Xiong, Z., Wang, D., Liu, X., Zhong, F., Wan, X., Li, X., Li, Z., Luo, X., Chen, K., Jiang, H., et al. Pushing the boundaries of molecular representation for drug discovery with the graph attention mechanism. Journal of medicinal chemistry, 63(16):8749–8760, 2019.

Yang, K., Swanson, K., Jin, W., Coley, C., Eiden, P., Gao, H., Guzman-Perez, A., Hopper, T., Kelley, B., Mathea, M., et al. Analyzing learned molecular representations for property prediction. Journal of chemical information and modeling, 59(8):3370–3388, 2019.

Yu, Q., Zhang, Y., Ni, Y., Feng, S., Lan, Y., Zhou, H., and Liu, J. Unified molecular modeling via modality blending. arXiv preprint arXiv:2307.06235, 2023.

Zhang, Z., Zhao, B., Xie, A., Bian, Y., and Zhou, S. Activity cliff prediction: Dataset and benchmark. arXiv preprint arXiv:2302.07541, 2023.

Zhou, G., Gao, Z., Ding, Q., Zheng, H., Xu, H., Wei, Z., Zhang, L., and Ke, G. Uni-mol: a universal 3d molecular representation learning framework. 2023.

